# Assessment of the Clinical Utility of Plasma Metagenomic Next-Generation Sequencing in a Pediatric Hospital Population

**DOI:** 10.1101/2020.03.06.981720

**Authors:** Rose A. Lee, Fatima Al Dhaheri, Nira R. Pollock, Tanvi S. Sharma

## Abstract

**Background:** Metagenomic next-generation sequencing (mNGS) of plasma cell-free DNA (cfDNA) is commercially available, but its role in the workup of infectious diseases is unclear.

**Methods:** To understand the clinical utility of plasma mNGS, we retrospectively reviewed patients tested at a pediatric institution over 2 years to evaluate the clinical relevance of the organism(s) identified and impact on antimicrobial management. We also investigated the effect of pre-test antimicrobials and interpretation of molecules of microbial cfDNA per microliter (MPM) plasma.

**Results:** 29/59 (49%) mNGS tests detected organism(s), and 28/51 (55%) organisms detected were clinically relevant. Median MPM of clinically relevant organisms was 1533 versus 221 for irrelevant organisms (p=0.01). mNGS test sensitivity and specificity were 53% and 79%, respectively, with a positive predictive value (PPV) of 72% and negative predictive value (NPV) of 50%. 14% of tests impacted clinical management by changing antimicrobial therapy. Immunocompromised status was the only patient characteristic that trended towards a significant clinical impact (p=0.056). No patients with culture-negative endocarditis had organisms identified by mNGS. There were no significant differences in antimicrobial pre-test duration between tests with clinically relevant organism(s) versus those that returned negative, nor was the MPM different between pre-treated and un-treated organisms, suggesting that 10 days of antimicrobial therapy as observed in this cohort did not sterilize testing; however, no pre-treated organisms identified resulted in a new diagnosis impacting clinical management

**Conclusions:** Plasma mNGS demonstrated higher utility for immunocompromised patients, but given the low PPV and NPV, cautious interpretation and Infectious Diseases consultation are prudent.

**Summary:** We evaluate the test performance characteristics and clinical utility of plasma metagenomic next-generation sequencing in a pediatric hospital cohort and demonstrate sensitivity and specificity of 53% and 79%, with 14% of tests impacting antimicrobial management.

## Introduction

Next-generation sequencing (NGS) describes high-throughput sequencing methods in which millions of DNA fragments can be independently and simultaneously sequenced. Cell-free DNA (cfDNA) in the bloodstream was first described in 1948^1^. CfDNA primarily originates from apoptotic human cells; inflammation, autoimmune disease, trauma, and cancer increase cfDNA levels^2-3^. NGS of cfDNA has been previously described for noninvasive diagnosis of fetal abnormalities^4-6^, cancer monitoring^7-10^, and transplant rejection^11-15^. Its adoption in these fields raised the prospect of diagnosing infections through sequencing of microbial cfDNA by metagenomic NGS (mNGS) followed by bioinformatic taxonomic classification.

mNGS, sometimes called shotgun sequencing, has been applied to various clinical sample types including cerebrospinal fluid, blood, respiratory samples, gastrointestinal fluid, and ocular fluid^16^. mNGS testing is “hypothesis-free,” unlike many contemporary molecular diagnostic infectious disease tests. Potential strengths include the ability to diagnose polymicrobial infections and quantitative reporting of cfDNA molecules detected. As blood traverses the entire body, it is hypothesized that even protected sites of infection may shed enough pathogen nucleic acid into blood for detection^17^. This pathogen-agnostic method is in contrast to targeted nucleic acid amplification tests (NAAT) that use specific primers, limiting detection to suspected targets. Because the vast majority of mNGS cfDNA reads will reflect the human host, sample processing methods for human DNA depletion are needed, supplemented by post-processing bioinformatic removal. Due to the amplification of background human DNA, mNGS is generally less sensitive than targeted approaches and requires greater sequencing depth for organism identification^18-19^.

A commercially available plasma cfDNA mNGS test from Karius Inc., (Redwood City, CA), available since 2016, reports molecules of microbial cfDNA per microliter (MPM) plasma. This laboratory is certified under the Clinical Laboratory Improvement Amendments of 1988, although the test has not been approved by the Federal Drug Administration. A recent company publication describes clinical and analytical test validation for detection of 1250 human pathogens^20^. The limit of detection of the Karius test is 41 MPM and organisms are reported if cfDNA from the organism is detected at statistically significant levels relative to negative controls run in parallel. For all reported organisms, a reference interval (MPM) is provided, based on abundances seen in samples from asymptomatic adult controls^20^. The relationship between MPM and microbe concentrations in blood [e.g. colony-forming units (CFUs)] is not well understood. Publications have described ongoing MPM detection for weeks after clearance of the organism on blood culture while on appropriate antimicrobial therapy^21^.

Despite potential strengths of cfDNA detection by mNGS, notable limitations exist. One obvious limitation is that the test will not detect RNA viruses. Importantly, uncertainty remains regarding how to assess if detected organism DNA (DNAemia) indicates a pathogen contributing to patient disease versus sample contamination or transient bacteremia from colonizing flora. In the clinical validation study by Karius Inc. ^20^, 350 patients who presented with sepsis alert criteria were tested and diagnostic sensitivity of 92.9% and specificity of 62.7% were reported in comparison to a composite reference standard, including all microbiological data and clinical history^20^. Sensitivity was 84.8% in comparison to standard microbiological testing alone. A recent study of 100 plasma mNGS tests sent from a pediatric hospital determined a sensitivity and specificity of the test for detection of organisms that impacted clinical decision-making of 92% and 64% respectively^22^.

At our hospital, clinicians have postulated that plasma mNGS may be useful in the following clinical scenarios: 1) culture-negative infections due to antibiotic pretreatment and/or fastidious or non-culturable organisms, and 2) deep-seated and difficult-to-sample infections such as invasive fungal infections, pneumonia, or osteomyelitis. The purpose of this study was to assess test performance characteristics and explore how mNGS findings impacted clinical management.

## Methods

We retrospectively reviewed medical records of all patients for whom commercial plasma mNGS testing was sent at Boston Children’s Hospital from October 2017 through October 2019. This study was approved by our institutional review board. Tests required approval from the directors of the Infectious Diseases (ID) Diagnostic Laboratory as well as an ID clinical consultation. The approval process involved a discussion about the utility of testing between the ID team and laboratory director when the diagnosis was not evident from initial testing. There were no fixed criteria and this study was conducted to help inform institutional guideline development based on identification of patient subsets in which the test was found to be the most clinically impactful. We assessed patient demographics, underlying comorbidities, ordering team, site of infection, duration of antimicrobial use prior to test, final clinical diagnoses, and reported MPM if testing returned positive for any organism. Patients were classified as immunocompromised if they had an underlying immunodeficiency, malignancy on active chemotherapy, hematopoietic stem cell or solid organ transplant, or other conditions requiring immunosuppression.

Clinical relevance of organisms identified from plasma mNGS was assessed relative to final overall diagnosis (infection versus no infection). Presence of an infection was determined by the treating clinical team and incorporated the clinical presentation that prompted mNGS testing and all microbiologic testing performed (including mNGS findings). A subgroup of clinically relevant organisms was “confirmed positive” if they correlated with a non-mNGS microbiological result (e.g. PCR or culture); however, in some cases, the clinical team made diagnoses on the basis of clinical picture and mNGS findings (Table 1A). These definitions of infection are consistent with prior studies that have evaluated the performance characteristics of mNGS^22,23^. In the absence of a gold standard for this novel technology, our composite reference standard nonetheless reflects how clinicians interpreted and acted on results, and we surmise this is the most clinically meaningfully definition of “infection”. Clinical relevance and confirmed positives were determined by expert opinions of two pediatric ID physicians not involved in the patient’s care at time of testing (R.L. and F.A.) with a tie-breaker opinion of a third (T.S.) if discordant.

**Table 1A:** Scenarios for clinically relevant (true positive) and clinically irrelevant (false positive/negative) organisms. *Example clinical scenario: concern for contaminant from standard microbiological testing and negative plasma mNGS results are used to clinically confirm suspicion and antibiotics are de-escalated

| Clinically Relevant (True positive): | Clinically Irrelevant (False positive or negative): |
| --- | --- |
| **Confirmed positive** and primary etiology of illness:<br>E.g. Patient septic from *Enterococcus* bacteremia on blood culture, which was also identified on mNGS testing | Pathogens that are likely **contaminant**: E.g. *Staphylococcus epidermidis* identified on mNGS but no evidence of bloodstream infection and concurrent blood cultures negative with no treatment |
| **Confirmed positive** but not primary reason for hospitalization/severe acute illness:<br>E.g. HSV gingivostomatitis in patient septic from *Pseudomonas* bacteremia, but HSV (and *Pseudomonas*) identified on mNGS testing and verified by standard workup (PCR swab and blood culture respectively) | Pathogens that may reflect **GI/skin colonization** with no obvious manifestation in the patient: e.g. *Neisseria sicca* co-identified in patient with respiratory failure/sepsis from adenovirus, and not confirmed on blood culture nor treated<br>Pathogens with **no known clinical significance**: e.g. virus with no known associated infectious clinical manifestation. |
| **Not Confirmed Positive but Consistent with Infectious Diagnosis**:<br>E.g. *Fusobacterium necrophilum* identified in mNGS testing in patient diagnosed with aspiration pneumonia, although standard microbiological workup didn't identify this organism | Pathogens identified on mNGS that were **discordant** with final clinical diagnosis made on the basis of standard microbiological workup: e.g. *Escherichia coli* and *Haemophilus influenzae* on mNGS in setting of *Streptococcus gordonii* endocarditis identified from blood culture and universal PCR of valve. |

**Table 1B:** Possible scenarios for determining clinical impact

| Plasma mNGS Result | Standard Microbiological Testing | Antimicrobial Change due to mNGS Result | Clinical Impact |  |
| --- | --- | --- | --- | --- |
| - | - | - | Redundant information, antibiotics and clinical plan were not changed (no impact) | - |
| - | - | + | Clinical impact (e.g. de-escalation) if team used negative mNGS results to de-escalate | + |
| - | + | - | No additional information (no impact) | - |
| - | + | + | Clinical impact (e.g. de-escalation) * | + |
| + | - | - | Not relevant organism (considered contamination or transient unrelated bacteremia) | - |
| + | - | + | Clinical impact (e.g. new diagnosis and targeted therapy) | + |
| + | + | - | Redundant information, antibiotics and clinical plan were not changed (e.g. known bacteria identified and no impact) | - |
| + | + | + | Clinical impact (e.g. different diagnosis and additional therapy) | + |

A novel aspect of our study was to assess the relationship of MPM to determination of a clinically relevant organism. We additionally considered whether there was antimicrobial use active against the organism by reviewing susceptibility data obtained via concurrent routine microbiological methods, when possible, and by assessing whether the patient clinically improved on empiric therapy, suggesting that it was appropriate.

We further evaluated the effect of mNGS testing on overall patient care to specifically assess the added value of plasma mNGS testing over standard microbiological workup, and defined “clinical impact” if testing resulted in 1) new organism(s) with new targeted antimicrobial therapy, 2) new organism(s) with de-escalation of antibiotic therapy, or 3) negative testing thus motivating teams to de-escalate antimicrobial therapy. Cases in which redundant organisms were identified on plasma mNGS and standard microbiological testing were only considered to have clinical impact if there was a change in antimicrobial management on the basis of the plasma mNGS result. For example, if the mNGS resulted in a diagnosis sooner than standard microbiological workup and affected antimicrobial management, this was considered to have a clinical impact. Clinical impact was adjudicated by the research team. Standard microbiological testing was defined as routine microbiological testing/NAAT performed either in our Infectious Diseases Diagnostic Laboratory or in reference laboratories. Logic gates of possible scenarios to determine clinical impact dependent on plasma mNGS, standard microbiological testing, and antimicrobial change are demonstrated in Table 1B.

### Statistical analysis

Demographic data were summarized using descriptive statistics. Test characteristics (sensitivity, specificity, negative and positive predictive value) for mNGS findings were calculated using two different methods (labeled as counting by test versus result) as illustrated in Figure 1 and Figure 2. Method 1 counted all mNGS results from one plasma sample as one test (n = 59). If the mNGS test sent identified a clinically relevant organism, whether or not the organism was a confirmed positive, the test result was considered a “true” positive. However, mNGS tests often identified multiple organisms, and in many of these instances, both clinically relevant and clinically irrelevant organisms (not related to any known or suspected infection in the patient) were reported. By method 1, the mNGS test would be classified as a true positive based on identification of a clinically relevant organism even if clinically irrelevant organism(s) were also identified. Method 1 therefore does not fully account for the “noise” of co-identified clinically irrelevant organisms. To account for this “noise”, we used Method 2 where we counted each organism identified so each organism result was assessed independently (n= 81). Method 2 provides more granular detail for mNGS findings by separately assessing the clinical relevance of each organism identified.

**Figure 1:**
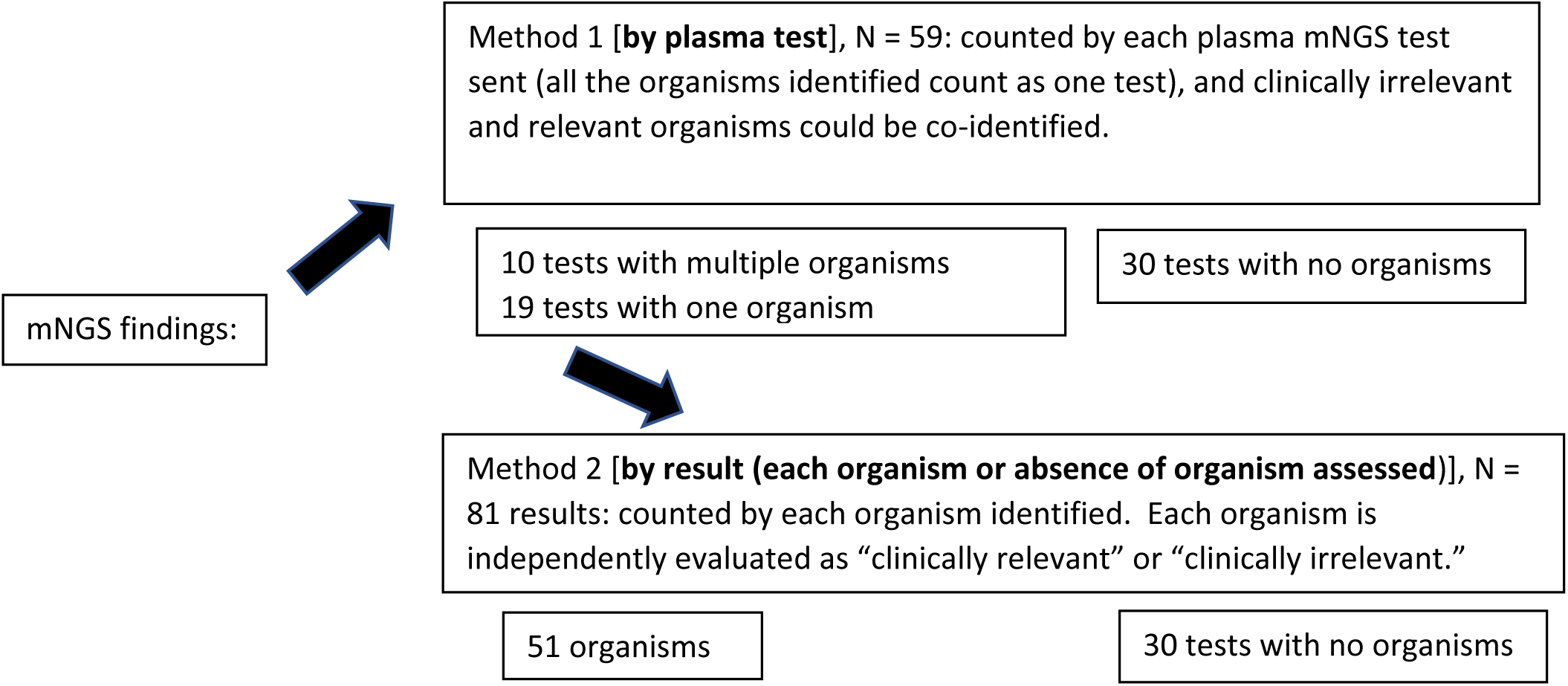
mNGS findings were counted by two separate methods, as illustrated above, for assessment of test characteristics by plasma test sent (Method 1), and by organism detected (Method 2).

**Figure 2:**
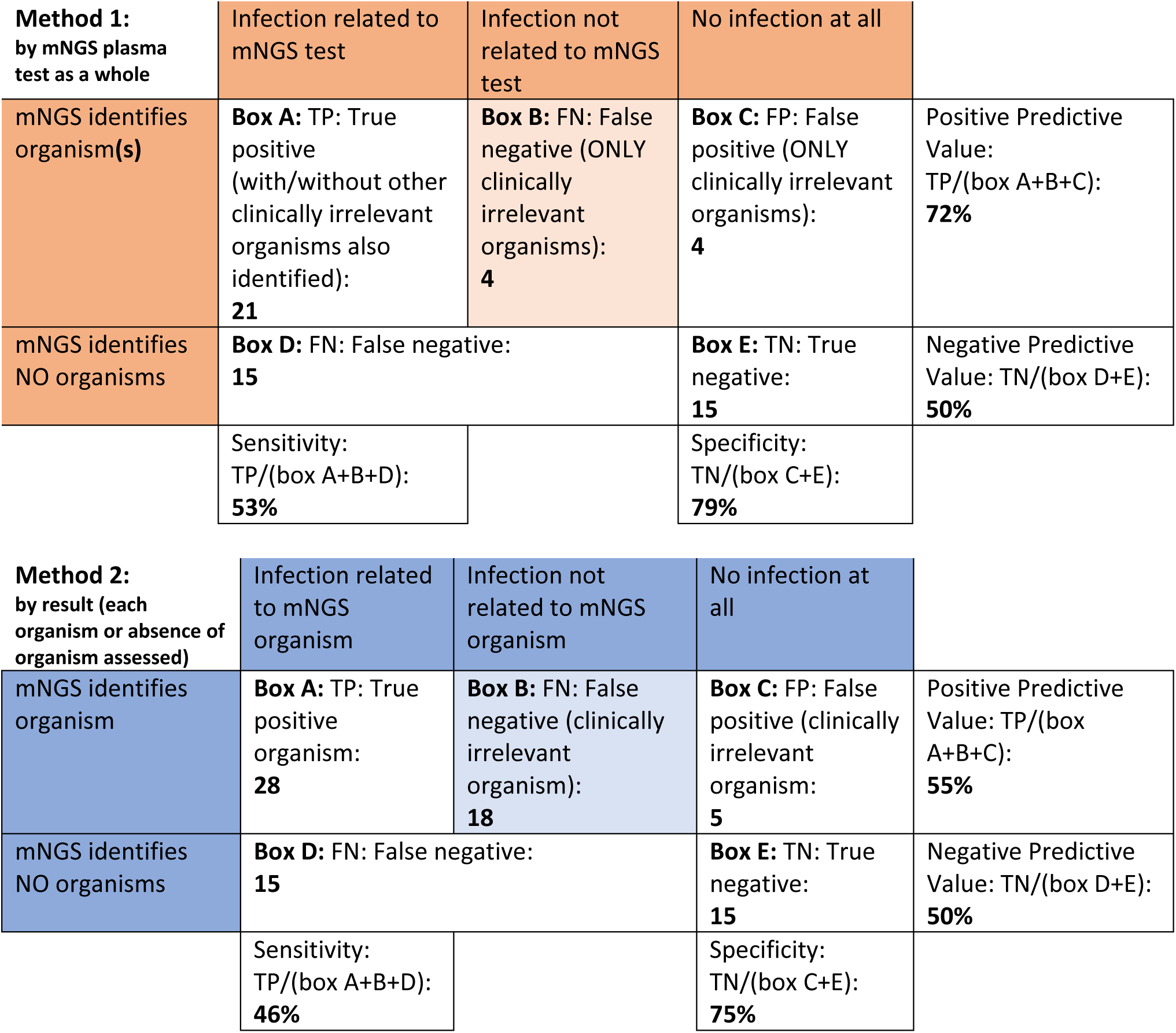
Testing characteristics calculated by Method 1 (each plasma test sent interpreted as a whole, n=59) and Method 2 (by organism) to discriminate noise in mNGS tests from clinically irrelevant organisms co-identified with relevant pathogens. Infection was defined by composite reference method (provider interpretation of clinical history and all microbiological data including mNGS findings). “Box B” was added to the usual 2×2 contingency table as these are clinically irrelevant organism(s) identified in the setting of an infection diagnosed by non-mNGS findings (i.e. diagnosed by standard microbiological workup). They cannot be included in Box D since mNGS identified organism(s) and cannot be included in Box C as the patient’s final diagnosis was infection. Nonetheless these cases contribute to sensitivity and positive predictive value and should not be dropped from calculations.

Comparative analysis was conducted by the Fisher’s exact test or chi-square test as appropriate and continuous data were compared using the Wilcoxon rank sums test and Kruskal-Wallis test for group medians. MPM performance in determination of clinically relevant organisms was assessed by receiver operating characteristics (ROC) analysis and area under the curve (AUC). An optimal cutoff score was found using the Youden index. Statistical tests were performed using Stata 15.1 software (Stata Corporation, College Station, TX, USA) and GraphPad v.8 software (GraphPad Software, San Diego, CA, USA) with p-values ≤ 0.05 as the significance threshold.

## Results

A total of 59 plasma NGS tests were sent on 54 patients during the study period. Table 2 summarizes patient characteristics, ordering teams, primary sites of infection, and final diagnoses of patients. Of the 5 tests that were re-sent on patients, two revealed new diagnoses (one with clinical impact) and all tests were sent at least a month apart with new or worsening clinical symptoms. The most common final diagnoses of patients on whom plasma mNGS was sent was no clear diagnosis (e.g. prolonged fever that could be due to infection or drug fever, but resolved without determination of specific etiology; 25%). Half of these patients were thought to ultimately have no infection at all, while the others were treated empirically for presumed infection. Autoimmune conditions were identified in 17% of patients and endocarditis in 14%. While cardiology teams ordered the second largest number of tests, no organisms were identified via mNGS on any of the culture-negative endocarditis cases and redundant organisms were identified in three cases by standard microbiological workup. In one case of culture-positive endocarditis, plasma mNGS identified discordant organisms that were deemed clinically irrelevant; *E.coli* and *H. influenzae* were identified on plasma mNGS but PCR of the eventually explanted valve identified *Streptococcus gordonii*, which also grew from an initial blood culture and was preliminarily considered a possible contaminant. No ordering team, primary site of infection, underlying comorbidities, or final patient diagnosis was noted to have a statistically significant association with clinical impact.

**Table 2:**
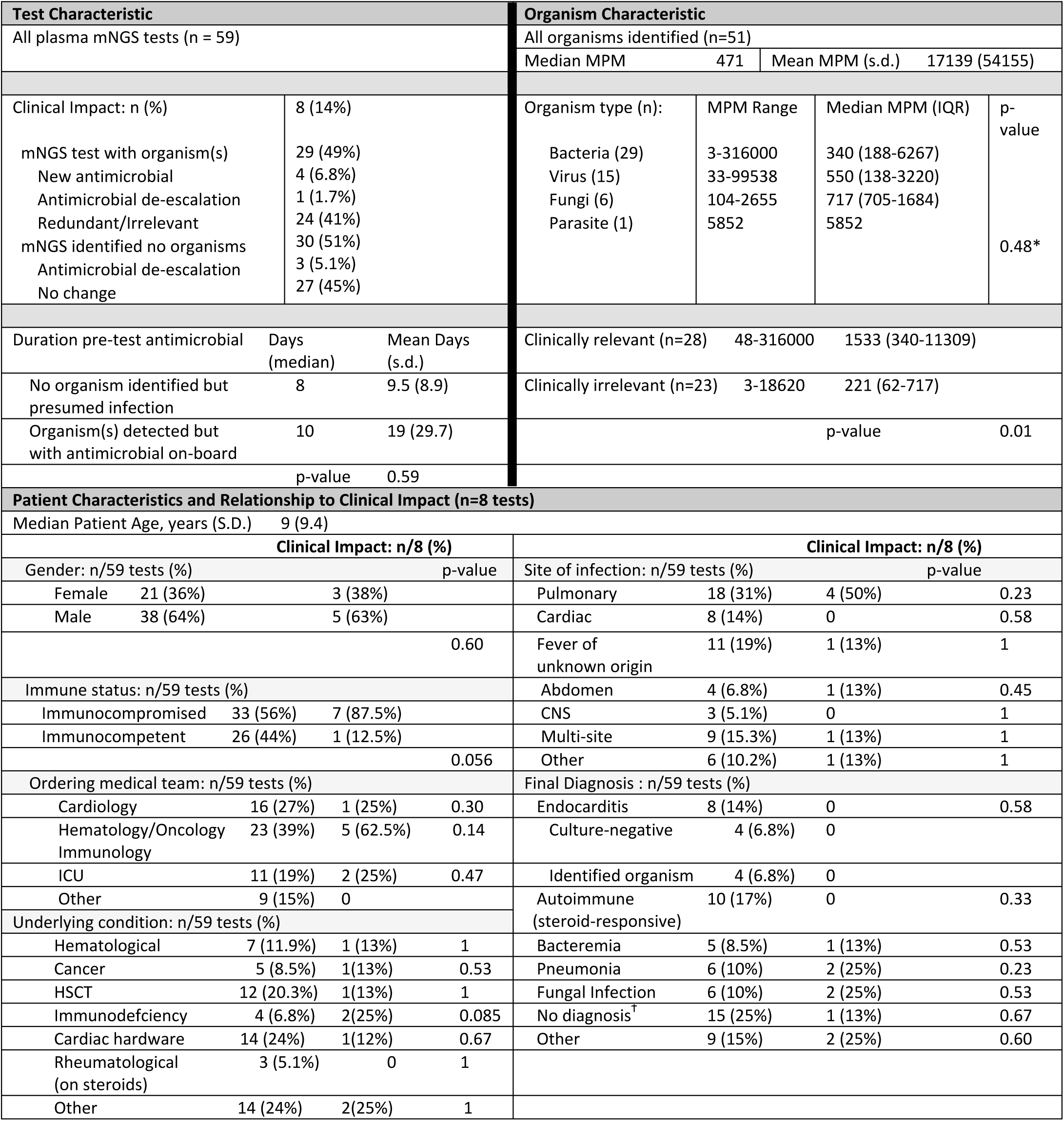
Plasma mNGS Test and Organism Characteristics, Clinical Impact, and Relevance. Patient characteristic p-values assess association of dichotomized categorical variable versus clinical impact by Fisher’s exact tests. *p-value to compare MPM medians by organism type did not include “parasite” as there was only one case. ^Ϯ^No diagnosis refers to no clear final diagnosis assigned by providers: 7 received empiric antimicrobials (assigned as infection), and 8 were ultimately considered to have no infection (no empiric antimicrobials)

Fifty-one organisms were identified from all testing combined (29 bacteria, 15 DNA viruses, 7 fungi, 1 parasite), 55% of which were considered clinically relevant. Table 2 summarizes the proportion of organisms identified that resulted in clinical impact or were determined to be redundant or clinically irrelevant.

In eight cases, testing led to clinical impact with a change (addition or de-escalation) in antimicrobial therapy. Seven out of the eight cases were immunocompromised patients and all of the five mNGS cases where a new organism was identified and new diagnosis was made impacting clinical management were in immunocompromised hosts (described in Supplementary Figure 1). Underlying immunodeficiency and overall immunocompromised status were the only variables found to trend towards a significant clinical impact although they did not reach our statistical threshold of 0.05 (p=0.08 and 0.06 respectively). While unexpected false positive and negative test results could lead to unnecessary investigations or treatment, we did not observe this in our cohort.

The sensitivity and specificity of plasma mNGS by test sent (method 1, n = 59) were 53% and 79%, respectively, with a positive predictive value (PPV) of 72% and negative predictive value (NPV) of 50% (Figure 2). Eight mNGS tests (14%) identified only clinically irrelevant organisms, and five mNGS tests deemed clinically relevant co-identified irrelevant organisms. When each organism identified was analyzed independently (method 2, n = 81), sensitivity/specificity were 46/75% with a PPV of 55% and NPV of 50% (Figure 2; organism and test assignments are described in Supplementary Dataset 1).

Testing was collected after a median of 8 days into clinical workup and median of 9 days of antimicrobial therapy, with median turnaround time (from time of receipt of sample by testing laboratory, to report) of 1 day, which is clinically actionable. For patients with plasma mNGS testing that returned negative in the setting of presumed infection treated empirically (“possibly sterilized” tests, n=15), antimicrobial therapy had been administered for a median of 8 days (mean 9.5, standard deviation 8.9) prior to test collection. Surprisingly, we found that the duration of pre-test therapy for patients with organisms detected on mNGS that should have been sterilized by the antimicrobial(s) in use (n = 27 organisms), was similar [median 10 days of therapy (p-value 0.59); mean 19, standard deviation 30]. For cases of presumed infection where both plasma mNGS and standard microbiological workup were negative, the majority of these infections were deep-seated infections (4 pulmonary infections, 2 osteomyelitis, 1 septic arthritis, 2 intrabdominal, 1 sepsis); four patients were diagnosed with culture-negative endocarditis.

We also assessed the relationship of MPM to identification of a clinically relevant organism. The median MPM for clinically relevant organisms was 1533 [interquartile range (IQR) 340-11309] in contrast to clinically irrelevant organisms (median MPM 221; IQR 62-717), which was a statistically significant difference (p=0.01). The median MPM for organisms with no pre-test antimicrobial therapy active against the organism was 407 (IQR 68-5852), compared to organisms with a covering antimicrobial (MPM 527; IQR 215-6267), which was not a statistically significant difference (p=0.78). While median MPMs did vary by organism type (Table 2), differences were not statistically significant (p=0.48 for bacteria versus fungi versus virus). A ROC curve for MPM data for distinction between clinically relevant and irrelevant organisms yielded an AUC of 0.75 (95% CI 0.611 to 0.887). An optimal cutoff of 390 MPM by Youden index was 74% sensitive (95% CI 55%-87%), and 73% specific (95% CI 52%-87%) with a likelihood ratio of 2.7 (Figure 3).

**Figure 3:**
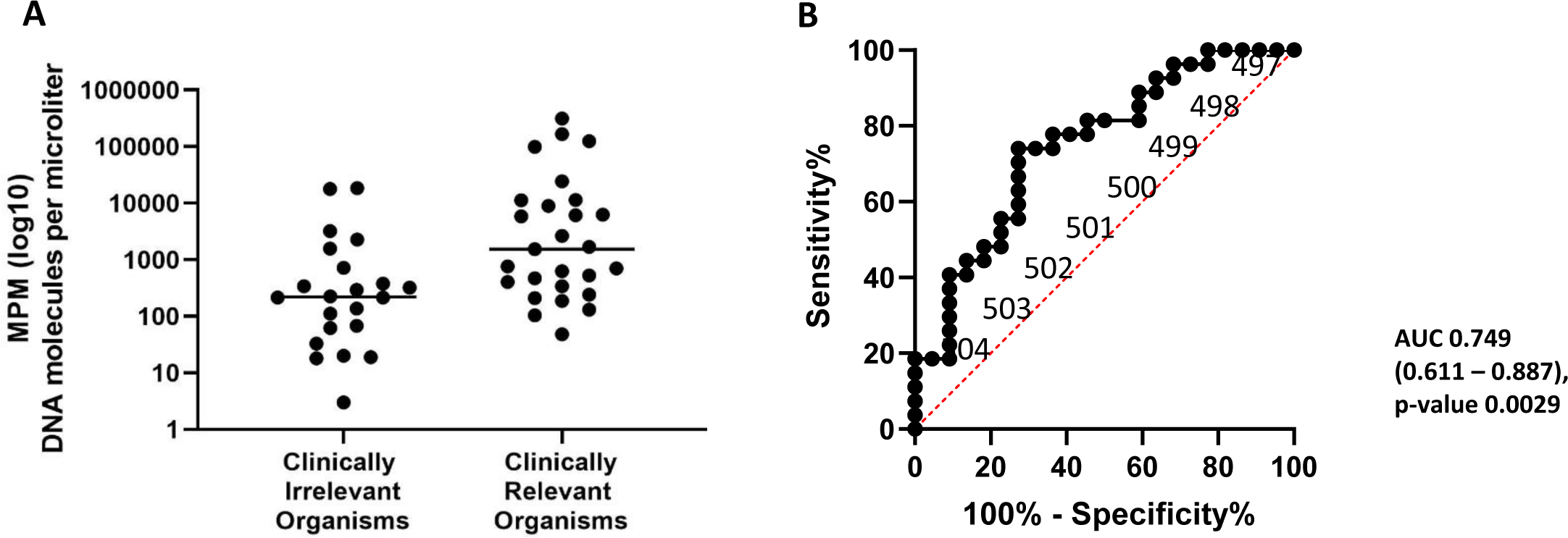
A: Comparison of distribution of MPM results for clinically relevant and irrelevant organisms (lines indicate medians) and B: Analysis of performance of MPM for distinction between clinically relevant and irrelevant organisms by receiver operating characteristic (ROC) curve.

## Discussion

In this study we describe the clinical utilization of plasma mNGS testing at our clinical center and include novel assessments not described in other studies. The sensitivity, specificity, PPV, and NPV of plasma mNGS testing at our hospital were considerably lower than results reported in the main clinical validation study led by the company^20^ as well as in a recent retrospective description of another pediatric hospital experience^22^. We surmise that the difference in test performance in part reflects a difference in how mNGS was applied, which was as a tertiary-level test sent in high-stakes scenarios where standard workup was unrevealing. At our institution, due to the considerable cost and unknown clinical utility, mNGS requires approval from the Infectious Diseases Diagnostic Laboratory Director and an ID consultation. We feel that our utilization likely reflects how many clinical centers would use plasma mNGS, in contrast to how this test was validated commercially as a sepsis screen in the emergency department^20^. This is the first study to account for the “noise” of polymicrobial identification in plasma mNGS in assessment of test performance and to individually assess the clinical relevance of each organism, which substantially impacted the positive predictive value (72% for per-test assessment versus 55% for per-organism assessment). We also included patients with a discordant mNGS finding (where the final clinical diagnosis of infection was made from standard microbiological workup and was not consistent with the mNGS finding) as cases for our calculations, rather than excluding them, in order to provide the most realistic estimates of test performance. Our study uniquely defined additional clinical factors we hypothesized could be relevant to plasma mNGS yield, including days into disease course, pre-test antimicrobial duration, and MPM interpretation.

This study illustrates how pretest probability affects testing utility, as the likelihood of plasma mNGS revealing an as-of-yet unidentified organism and new diagnosis after standard workup was low, particularly for immunocompetent patients. Many of our patients ultimately had a non-infectious diagnosis, or a presumed infection treated empirically in the absence of microbiological data, which yielded higher false positives and negatives in comparison to prior studies. Negative mNGS results in patients with culture-negative infections (designated as false negatives) also mostly involved protected sites of infection (pulmonary, intrabdominal, bone), which is suggestive that plasma mNGS may be an inadequate and at worst a misleading proxy for invasive microbiological sampling. Notably, the test had minimal yield for culture-negative endocarditis, despite the adjacency of cardiac valves to blood (only one endocarditis case underwent surgical management and had confirmed endocarditis on pathology, but all cases had presentations that met modified Duke’s criteria for endocarditis and improved on therapy). We additionally report that the clinical impact of tests through changes in antimicrobial therapy was low (14%), although notably this was higher than another study that found that only 7% of tests led to a positive clinical impact^21^.

A key overall finding was that the negative predictive value in our clinical practice was only 50%. While many providers wanted to use plasma mNGS to “rule-out” an infection, we show that negative tests only predict the absence of an infection as well as a coin flip, and therefore are a poor rule-out screening test. However, we did find a significant association between MPM reported and clinical relevance (Figure 3), suggesting that high MPMs should make providers more confident that the result is meaningful.

Given that mNGS was sent several days into the disease course, we also wanted to address the possible impact of empiric pre-test antimicrobials on plasma mNGS yield. While clearance of bloodstream pathogen cfDNA over time is expected, kinetics for specific pathogens will need to be elucidated as mNGS becomes more routine. Counterintuitively, we did not find significant differences in MPM values between organisms treated with an appropriate antimicrobial pre-test and those untreated, even when only considering clinically relevant organisms (dismissing organisms that may have been contaminants and thus unaffected by antimicrobials). Furthermore, we did not find significant differences in antimicrobial duration between “possibly sterilized” mNGS tests and tests where an organism was identified with an active antimicrobial on-board. This suggests that pre-test antimicrobial durations of 10 days (median) as observed in this cohort do not likely substantially affect sterilization of plasma mNGS. The ongoing detectable MPM may be related to slow-to-clear DNAemia from high pathogen burden even though organisms may have been appropriately killed on targeted therapy, a finding that is consistent with prior reports.^22^ Notably, no identified pre-treated organisms resulted in a novel diagnosis that affected clinical management in our cohort.

Limitations of this study include a relatively small sample size, which in turn leads to a small number of patients in each relevant diagnostic sub-category (e.g. culture-negative endocarditis) and for establishment of the MPM cutoff in ROC analysis. Additionally, our gold standard definition of the presence of infection was a composite assessment from the provider team, which included interpretation of all microbiological data including mNGS findings. In the ideal scenario, we would have an independent gold standard of the test under evaluation although there is precedent in the literature for assessing novel and possibly more sensitive technologies this way^23-25^. In clinical practice, providers routinely incorporate the results of this test with other clinical data and, understanding the limitation that there is no reference standard for mNGS, our goal was to characterize provider response to findings, in the context of all of the information available for the patient.

In summary, our major findings included lower sensitivity and specificity of plasma mNGS than prior literature suggests, with only half of the organisms identified as clinically relevant -- emphasizing the need for ID consultation for interpretation. We found higher utility for immunocompromised patients, and less value than expected for endocarditis. Additionally, although we expected that pre-test antimicrobials would decrease the yield of plasma mNGS testing, after 10 days (median) of antimicrobial therapy, the MPM did not differ significantly between treated and untreated organisms nor was overall detection compromised. Despite the insights gained in this study regarding plasma mNGS test performance and utility, further work will be required to understand how to optimally integrate this technology into the infectious diseases diagnostic work up.

## Acknowledgements

We thank K.P. Smith for his insightful review and comments on this manuscript.

## References

1) Mandel P. 1947. Les acides nucleiques du plasma sanguin chez 1 homme. CR Seances Soc Biol Fil 142: 241–3.

2) Leon SA, Shapiro B, Sklaroff DM, Yaros MJ. 1977. Free DNA in the serum of cancer patients and the effect of therapy. Cancer Res 37(3): 646–50.

3) Lo YD, Rainer TH, Chan LY, Hjelm NM, Cocks RA. 2000. Plasma DNA as a prognostic marker in trauma patients. Clin Chem 46(3): 319–23.

4) Stokowski R, Wang E, White K, Batey A, Jacobsson B, Brar H, Balanarasimha M, Hollemon D, Sparks A, Nicolaides K, Musci TJ. 2015. Clinical performance of non-invasive prenatal testing (NIPT) using targeted cell-free DNA analysis in maternal plasma with microarrays or next generation sequencing (NGS) is consistent across multiple controlled clinical studies. Prenat Diagn 35: 1243–1246.

5) Song, K, Musci TJ, Caughey AB. 2013. Clinical utility and cost of noninvasive prenatal testing with cfDNA analysis in high-risk women based on a US population. J Matern Fetal Neonat Med 26: 1180–1185.

6) Fan HC, Blumenfeld YJ, Chitkara U, Hudgins L, Quake SR. 2008. Noninvasive diagnosis of fetal aneuploidy by shotgun sequencing DNA from maternal blood. Proc Natl Acad Sci USA 105: 16266–16271.

7) Aravanis AM, Lee M, Klausner RD. 2017. Next-generation sequencing of circulating tumor DNA for early cancer detection. Cell 168: 571–574.

8) Lanman RB, Mortimer SA, Zill OA, Sebisanovic D, Lopez R, Blau S, Collisson EA, Divers SG, Hoon DS, Kopetz ES, Lee J. 2015. Analytical and clinical validation of a digital sequencing panel for quantitative, highly accurate evaluation of cell-free circulating tumor DNA. PLoS One 10:e0140712.

9) Bettegowda C, Sausen M, Leary RJ, Kinde I, Wang Y, Agrawal N, Bartlett BR, Wang H, Luber B, Alani RM, Antonarakis ES. 2014. Detection of circulating tumor DNA in early- and late-stage human malignancies. Sci Transl Med 6: 224ra224.

10) Dawson SJ, Tsui DW, Murtaza M, Biggs H, Rueda OM, Chin SF, Dunning MJ, Gale D, Forshew T, Mahler-Araujo B, Rajan S. 2013. Analysis of circulating tumor DNA to monitor metastatic breast cancer. N Engl J Med 368: 1199–1209.

11) Schütz E, Fischer A, Beck J, Harden M, Koch M, Wuensch T, Stockmann M, Nashan B, Kollmar O, Matthaei J, Kanzow P. 2017. Graft-derived cell-free DNA, a noninvasive early rejection and graft damage marker in liver transplantation: a prospective, observational, multicenter cohort study. PLoS Med 14: e1002286.

12) Bloom RD, Bromberg JS, Poggio ED, Bunnapradist S, Langone AJ, Sood P, Matas AJ, Mehta S, Mannon RB, Sharfuddin A, Fischbach B. 2017. Cell-free DNA and active rejection in kidney allografts. J Am Soc Nephrol 28: 2221–2232.

13) De Vlaminck I, Martin L, Kertesz M, Patel K, Kowarsky M, Strehl C, Cohen G, Luikart H, Neff NF, Okamoto J, Nicolls MR. 2015. Noninvasive monitoring of infection and rejection after lung transplantation. Proc Natl Acad Sci USA 112: 13336–13341.

14) De Vlaminck I, Valantine HA, Snyder TM, Strehl C, Cohen G, Luikart H, Neff NF, Okamoto J, Bernstein D, Weisshaar D, Quake SR. 2014. Circulating cell-free DNA enables noninvasive diagnosis of heart transplant rejection. Sci Transl Med 6: 241ra277.

15) Snyder TM, Khush KK, Valantine HA, Quake SR. 2011. Universal noninvasive detection of solid organ transplant rejection. Proc Natl Acad Sci USA 108: 6229–6234.

16) Gu W, Miller S, Chiu C. 2019. Clinical Metagenomic Next-Generation Sequencing for Pathogen Detection. Annu Rev Pathol 14: 319–338.

17) Farnaes L, Wilke J, Loker KR, Bradley JS, Cannavino CR, Hong DK, Pong A, Foley J, Coufal NG. 2019. Community-acquired pneumonia in children: cell-free plasma sequencing for diagnosis and management. Diagn Microbiol Infect Dis 94(2): 188–91.

18) Cummings LA, Kurosawa K, Hoogestraat DR, SenGupta DJ, Candra F, Doyle M, Thielges S, Land TA, Rosenthal CA, Hoffman NG, Salipante SJ. 2016. Clinical next generation sequencing outperforms standard microbiological culture for characterizing polymicrobial samples. Clin Chem 62(11): 1465–73.

19) Salipante SJ, Kawashima T, Rosenthal C, Hoogestraat DR, Cummings LA, Sengupta DJ, Harkins TT, Cookson BT, Hoffman NG. 2014. Performance comparison of Illumina and ion torrent next-generation sequencing platforms for 16S rRNA-based bacterial community profiling. Appl Environ Microbiol 80(24): 7583–91.

20) Blauwkamp TA, Thair S, Rosen MJ, Blair L, Lindner MS, Vilfan ID, Kawli T, Christians FC, Venkatasubrahmanyam S, Wall GD, Cheung A. 2019. Analytical and clinical validation of a microbial cell-free DNA sequencing test for infectious disease. Nat Microbiol 4(4): 663.

21) Hogan CA, Yang S, Garner OB, Green DA, Gomez CA, Bard JD, Pinsky BA, Banaei N. 2020. Clinical Impact of Metagenomic Next-Generation Sequencing of Plasma Cell-Free DNA for the Diagnosis of Infectious Diseases: A Multicenter Retrospective Cohort Study. Clinical Infectious Diseases ciaa035.

22) Fung M, Zompi S, Seng H, Hollemon D, Parham A, Hong DK, Bercovici S, Dolan E, Lien K, Teraoka J, Logan AC. 2018. Plasma Cell–Free DNA Next-Generation Sequencing to Diagnose and Monitor Infections in Allogeneic Hematopoietic Stem Cell Transplant Patients. Open Forum Infect Dis 5(12): ofy301

23) Rossoff J, Chaudhury S, Soneji M, Patel SJ, Kwon S, Armstrong A, Muller WJ. 2019. Noninvasive Diagnosis of Infection Using Plasma Next-Generation Sequencing: A Single-Center Experience. Open Forum Infect Dis 6(8): ofz327.

24) Miao Q, Ma Y, Wang Q, Pan J, Zhang Y, Jin W, Yao Y, Su Y, Huang Y, Wang M, Li B. 2018. Microbiological diagnostic performance of metagenomic next-generation sequencing when applied to clinical practice. Clin Infect Dis 67(suppl_2):S231–40.

25) McAdam AJ. 2000. Discrepant analysis: how can we test a test? J Clin Microbiol. 38(6): 2027–9.

